# Deep learning-assisted single-molecule detection of protein post-translational modifications with a biological nanopore

**DOI:** 10.1101/2023.09.08.555904

**Authors:** Chan Cao, Pedro Magalhães, Lucien Fabrice Krapp, Juan F. Bada Juarez, Simon Mayer, Verena Rukes, Anass Chiki, Hilal A. Lashuel, Matteo Dal Peraro

**Affiliations:** Institute of Bioengineering, School of Life Sciences, Ecole Polytechnique Fédérale de Lausanne, EPFL, Lausanne 1015, Switzerland; Department of Inorganic and Analytical Chemistry, University of Geneva, 1211 Geneva, Switzerland; Laboratory of Molecular and Chemical Biology of Neurodegeneration, Brain Mind Institute, School of Life Sciences, Ecole Polytechnique Fédérale de Lausanne, EPFL, Lausanne 1015, Switzerland

**Keywords:** Biological nanopores, protein post-translational modifications, deep-learning, single-molecule sensing, α-synuclein

## Abstract

Protein post-translational modifications (PTMs) play a crucial role in countless biological processes, profoundly modulating protein properties on both the spatial and temporal scales. Protein PTMs have also emerged as reliable biomarkers for several diseases. However, only a handful of techniques are available to accurately measure their levels, capture their complexity at a single molecule level and characterize their multifaceted roles in health and disease. Nanopore sensing provides high sensitivity for the detection of low-abundance proteins, holding the potential to impact single-molecule proteomics and PTM detection in particular. Here, we demonstrate the ability of a biological nanopore, the pore-forming toxin aerolysin, to detect and distinguish α-synuclein-derived peptides bearing single or multiple PTMs, namely phosphorylation, nitration and oxidation occurring at different positions and in various combinations. The characteristic current signatures of the α-synuclein peptide and its PTM variants could be confidently identified using a deep learning model for signal processing. We further demonstrate that this framework can quantify α-synuclein peptides at picomolar concentration and detect the C-terminal peptides generated by digestion of full-length α-synuclein. Collectively, our work highlights the unique advantage of using nanopore as a tool for simultaneous detection of multiple PTMs and paves the way for their use in biomarker discovery and diagnostics.

## Introduction

Proteins are the major molecular building blocks of life, dictating the spatial and temporal occurrence of most biological functions^1^. Their functional properties are mainly determined by their native three-dimensional structure, subcellular localization and interactome. Post-translational modifications (PTMs) play an important role in regulating all these properties of proteins and thus represent molecular switches for coordinating, diversifying and regulating cellular function^2^. Moreover, the majority of protein-based biopharmaceuticals approved or in clinical trials bear some form of PTMs in order to enhance their efficacy for therapeutic applications^3^. Despite the impressive recent advances in biochemistry and structural biology, the development of sensitive methods to detect PTMs, generate site-specifically modified proteins, and decipher the PTM code of proteins still lags behind.

The standard methods currently available for the detection of PTMs are mass spectrometry (MS), and enzyme-linked antibody-based assays^4,5^, while immunoprecipitation coupled to MS (IP/MS) has also been increasingly used to detect and quantify low abundant protein species. Recently, great progress has been made towards developing and improving MS methods and immunoassays to map, detect and quantify modified proteins in biological samples. For instance, proximity extension assays^6^ and single molecule array (Simoa)^7^ have shown high sensitivity for detecting PTMs. One of the major drawbacks of these immunoassays is that they are based on antibodies that are developed against single PTMs. The development and optimization of these assays did not account for the presence of multiple/neighbouring PTMs. One example is demonstrated in our recent work, which shows that neighboring PTMs interfere with the detection of phosphorylated α-synuclein at S129 by most pS129 antibodies^8^. Therefore, a method to accurately identify protein PTMs is essential as increasing evidence suggests the PTM code is combinatorial in nature and involves complex interplay and cross-talk among PTMs^9,10^.

PTMs have been implicated in a wide variety of diseases, such as characterizing the changes in protein PTM types and levels is highly relevant for early diagnosis and monitoring the progression of diseases, as well as for evaluating the efficacy of new therapies^4,11^. For example, several neurodegenerative diseases including Alzheimer’s disease (AD) and Parkinson’s disease (PD) are caused by the accumulation of misfolded and aggregated proteins in the brain regions that are affected by the disease (*e*.*g*., amyloid-β in amyloid plaques, Tau in neurofibrillary tangles and α-synuclein in Lewy bodies and Lewy neurites). One shared characteristic among these proteins is that their aggregated forms are heavily modified. PTMs have emerged as key signatures of disease pathologies and are commonly used as the primary (bio)markers of disease progression and pathology formation, spreading and clearance in response to therapies^12,13^.

Nanopore sensing is an approach based on ionic current readout that can detect a single molecule as it is passing through a nanometer scale pore^14^. Such pores are either biological assemblies of proteins embedded in a lipid membrane or are fabricated by a solid-state material^15^. When a molecule of interest passes through such a pore, the electric current signal is modulated and exquisitely sensitive to the molecule of interest and thus can provide information about its size, mass, charge, composition, structure and conformation in real-time^16,17^. Nanopore sensing has achieved sequencing of ultra-long DNA^18,19^, and this success has inspired its application for peptides and proteins analysis^20– 23^. Compared with MS, nanopore sensing is faster, cheaper, free of ionization and has an ultrasensitive detection of biomolecules in solution at the single-molecule level. These unique features, in particular its sensitivity, make nanopores perfectly suited for protein analysis since there is no biochemical method available for protein amplification, like polymerase chain reaction that has been widely used for DNA amplification. Recently, several groups have explored the potential of using nanopore approaches to detect protein PTMs, including phosphorylation^24–26^, acetylation^27^, propionylation^28^, glycosylation^29,30^ and ubiquitination^31^. However, these studies either focus on identifying different types of modifications or the same modification at different positions, but do not consider the simultaneous occurrence of different types and combinations of PTM species that are of clear clinical relevance. Moreover, the detection of low abundant protein PTMs from clinical samples is still poorly investigated.

Here, we demonstrate that an engineered form of aerolysin^32–35^, can be used to detect peptides containing single or multiple types of PTMs. As a model system, we used peptides derived from the C-terminal domain of the presynaptic protein α-synuclein, the primary constituent of the pathological hallmarks of PD (*i*.*e*., Lewy bodies and Lewy neurites). This region is characterized by the clustering of different types of PTMs (*e*.*g*., phosphorylation, nitration and oxidation) that are associated with pathology formation and disease progression in PD and other neurodegenerative diseases^36^. Recent studies also suggest that specific α-synuclein PTMs (*i*.*e*., pS129) are elevated in the cerebral spinal fluid (CSF) of PD patients and could be developed into diagnostic biomarkers for PD and potentially for other neurodegenerative diseases^37,38^.

We observe that phosphorylation, nitration and oxidation significantly modulate the current amplitude and dwell time of the signals in comparison with the wild-type (wt) α-synuclein. Additionally, we show that the α-synuclein peptides bearing multiple PTMs and different PTMs combinations induce unique ionic current signatures, which opens the door to investigate PTMs crosstalk and monitor the PTMs dynamics in the future. A deep learning approach was specifically developed to process the signals, providing an automatic and fast way to classify current readouts associated with each PTM combination. We also demonstrate the ability of aerolysin pores to detect unmodified α-synuclein peptides when mixed with red blood cells (RBCs) at picomolar concentration, as well as the possibility of using cathepsin D to generate the specific C-terminal peptides that bear most PTMs, providing promising ground for actual clinical applications. In summary, our work presents a single-molecule approach able to detect clinically relevant, low abundance α-synuclein PTMs. Compared to other analytical methods like MS and ELISA^39,40^, this nanopore-based approach offers detection at single-molecule resolution and provides promising ground to decipher PTMs patterns relevant for disease diagnostics.

## Results

### Characterization of the C-terminal region of α-synuclein with a nanopore

The ability of aerolysin to detect PTMs was examined by means of single-channel recording experiments as shown in **Figure 1a**. A lipid membrane separates the chamber into two compartments, *cis* and *trans*, and holds an aerolysin pore that connects them. An engineered variant of aerolysin is used here, namely K238A, where lysine residues at position 238 are substituted by alanine residues, providing a significantly enhanced resolution for biomolecular sensing compared to wt aerolysin^41^. As mentioned above, we focused on α-synuclein as a model system because of the strong links between its PTM profile and the brain pathology of several neurodegenerative diseases, most notably in PD_36,42_.

**Figure 1.**
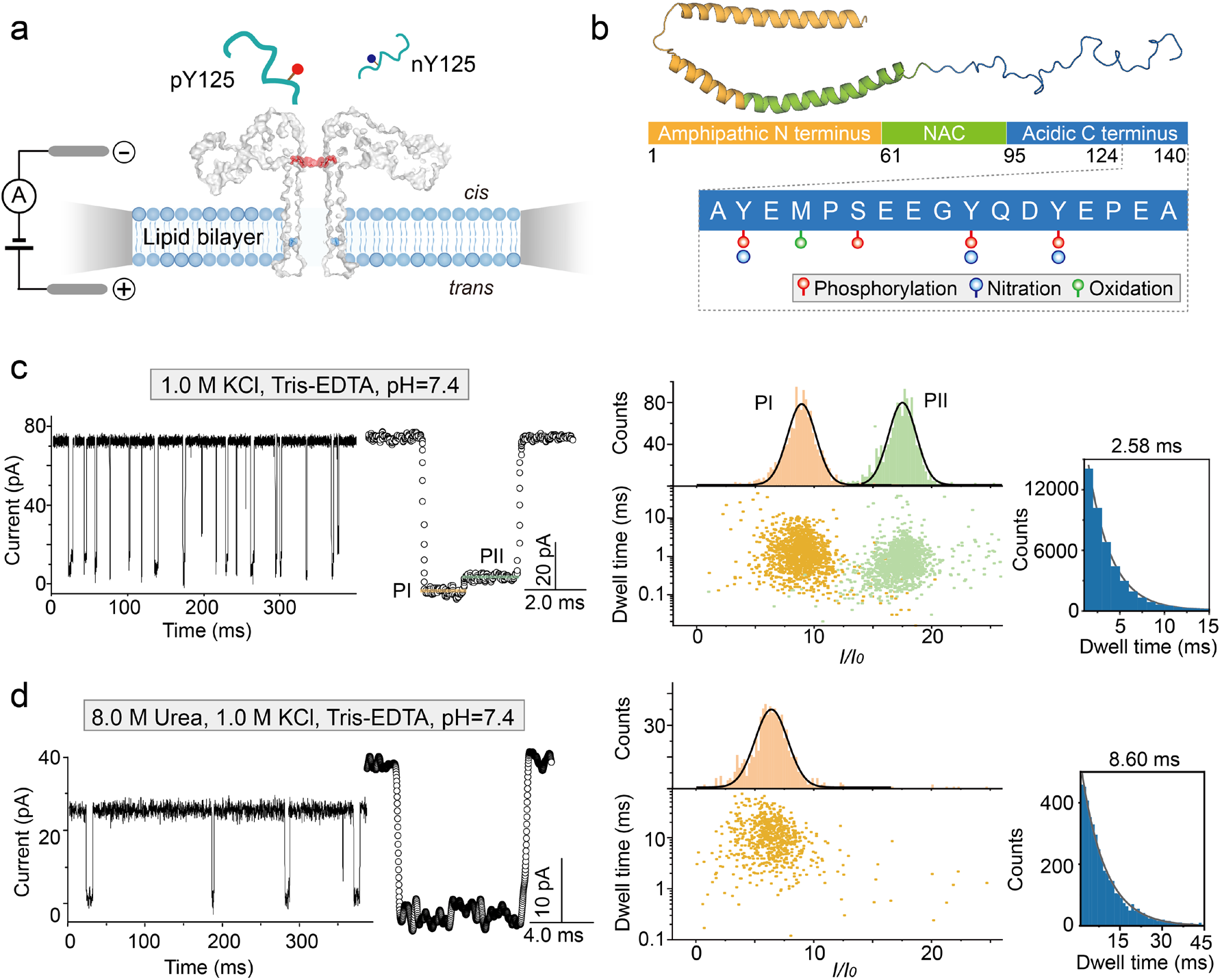
Characterization of wild-type α-synuclein C-terminal peptides. **(a)** Illustration of the single-channel recording setup composed of two chambers separated by a lipid membrane that is formed across an orifice; the chambers are named *cis* and *trans*. Voltage is applied across the pore using two Ag/AgCl electrodes. **(b)** The structure of α-synuclein in full-length (PDB:1xq8), including the amphipathic N-terminus (yellow), non-amyloid-component (NAC) domain (green), and the disordered C-terminal fragment (blue). Phosphorylation (red circle), nitration (blue circle) and oxidation (green circle) of residues 124-140 are highlighted. Nanopore single channel recording of wt α-syn_124-140_ peptide in 1.0 M KCl **(c)** and in 1.0 M KCl, 8.0 M urea **(d)** solutions, respectively, both buffered with 10 mM Tris, and 1.0 mM EDTA at pH 7.4. Left panels of **(c)** and **(d)** report raw current traces, middle panels typical events, while the right panels show scatter plots, *I/I*_*0*_ percentage and dwell time histograms. All data were obtained by applying a voltage of +100 mV.

The C-terminal domain of α-synuclein, encompassing the residues 124-140 (α-syn_124-140_) (**Figure 1b**) harbors several PTMs that are found in pathological α-synuclein aggregates in the brain of patients with PD and other neurodegenerative diseases^42^. Many of these PTMs, including phosphorylation and nitration, have emerged as reliable markers of α-synuclein pathology formation in human brains and animal models of PD and related synucleinopathies^36^. The C-terminal domain of α-synuclein is rich in proline residues, it is highly negatively charged and does not adopt a stable secondary structure in the monomeric state of the protein^43^, which in this context is particularly convenient as these peptides can be easily driven to translocate through the nanopore by the potential applied across the lipid membrane. All peptides were prepared using Fmoc-based solid-phase peptide synthesis and purified as previously reported^44–46^ (see Materials and Methods).

α-Syn_124-140_ was added into the *cis* compartment and when a positive voltage was applied to the *trans* compartment, clear and reproducible blockades of ionic current were obtained (**Figure 1c-d**). For wt α-syn_124-140_, a peculiar and well-recognizable 2-level blockade event was observed and the signal often contained a higher residual current at the last fraction (**Figure 1c**, middle). As a result, two populations were observed (**Figure 1c**, right): one exhibited a relative current percentage (*I/I*_*0*,_ see Materials and Methods) of 9.0 ± 2.0, while the other 17.1 ± 2.0 (named hereafter *PI* and *PII*, respectively). These values correspond to the mean and standard deviation derived from a Gaussian fit of the relative current histogram.

To gain further insights into this peculiar two-level signal, we considered the structure of this α-synuclein fragment. Based on previous studies^47^, there is a highly hydrophobic segment between residues 125-129 that can produce a structurally compact cluster likely responsible for the initial lower residual current. To test this hypothesis, we reasoned that destabilizing this conformation by the addition of urea could produce signals expected for a more extended peptide^47^. We thus used 8.0 M urea (along with 1.0 M KCl, 10 mM Tris, 1.0 mM EDTA, and pH 7.4) to induce complete unfolding of the peptide while aerolysin remained folded and functional (**Supplementary Figure 1**). Under these conditions, the open pore current was lower (25 ± 1.8 pA at 100 mV) compared to the same salt concentration without urea (72 ± 1.5 pA at 100 mV). This is because the additional urea significantly decreased the mobility of the ions^48,49^. However, in these conditions, wt α-syn_124-140_ translocated with a simple one-level signal as evidenced by one population in the scatter plot and the *I/I*_*0*_ histograms (**Figure 1d**). These one-level current signals suggest that the addition of urea results in a more extended conformation of the polypeptide chain, which facilitates its translocation through the pore. In addition, due to the decrease of ionic mobility, the dwell time became much longer, 8.6 ± 0.24 ms, approximately 4 times longer than the condition without urea. These results demonstrate that the conformation of largely disordered peptides like those obtained from α-synuclein can be monitored at the single-molecule level using a nanopore.

### Deep learning-assisted detection of different PTMs and their combinations

Next, we assessed the feasibility of using nanopore for detecting PTMs of α-syn_124-140_ peptide. We investigated different types of modifications occurring at the same amino acid (i.e., phosphorylation and nitration of Y125, pY125 and nY125), or the same type of PTM located at different positions (i.e., pS129 and pY125, nY125 and nY136). Moreover, we considered peptides bearing multiple PTMs of the same type (i.e., pY125pS129 and nY125nY133nY136) and of different types (i.e., nY125pS129).

As shown in **Figure 2a**, the current signatures of the different PTM combinations appeared all to have distinct current characteristics. Compared to the unmodified peptide, the pY125 and nY125 α-syn_124-140_ peptide presented only one level and their relative current were lower than the wt peptide (**Supplementary Figure 2**). This decrease in the relative current value is consistent with a deeper blockade induced by the additional volume due to the presence of the PTMs. Moreover, the dwell time varied significantly with different types of PTMs, the fitted values being 2.58±0.4 ms for wt, 0.55±0.08 ms for pY125 and 4.51±0.5 ms for nY125, indicating that phosphorylation speeds up the translocation process, while nitration significantly slows it down. The faster translocation of pY125 could be induced by the additional negatively charged phosphate group. In addition, we investigated the effect of oxidation of methionine at position 127 of ±-syn_124-140_ (oM127, **Supplementary Figure 3**). Similar to pY125 and nY125, oM127 also showed only one current level.

**Figure 2.**
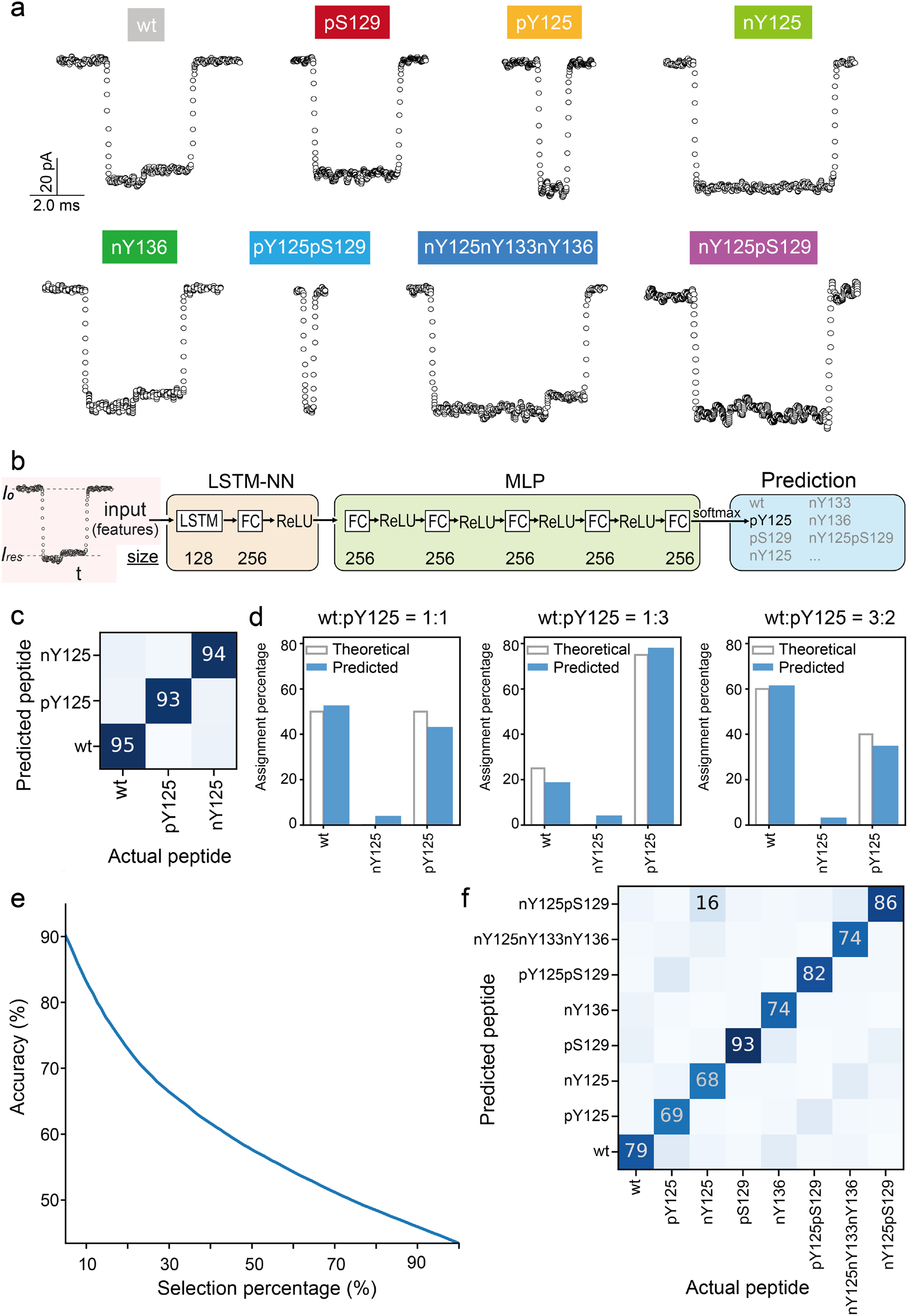
Discrimination of α-synuclein peptide PTMs by deep learning. **(a)** Typical ionic current signals of different PTM types and their spatial combinations. The scale shown in the left top panel applies to all the events. **(b)** Deep learning approach to signal processing: an LSTM recurrent neural network was used to read the events and then followed by an MLP to predict the peptides. **(c)** Confusion matrix of wt, pY125 and nY125 peptides obtained by deep learning approach. **(d)** Assignment percentage of different mixture samples of wt and pY125 at a concentration ratio of 1:1 (left), 1:3 (middle) and 3:2 (right). The theoretical accuracy is shown by the white columns while the predicted accuracy is represented by blue columns. **(e)** Selection percentage versus averaged accuracy obtained from deep learning approach of peptides, including wt, pY125, nY125, pS129, pY125pS129, nY125nY133nY136 and nY125pS129. **(f)** The confusion matrix of wt, pY125, nY125, pS129, pY125pS129, nY125nY133nY136 and nY125pS129classification. Columns represent actual peptides from the test set, while rows are the peptides that the deep learning algorithm assigned them to. All data were obtained using 1.0 M KCl, 10 mM Tris, and 1.0 mM EDTA buffer at pH 7.4 applying a voltage of +100 mV.

**Figure 3.**
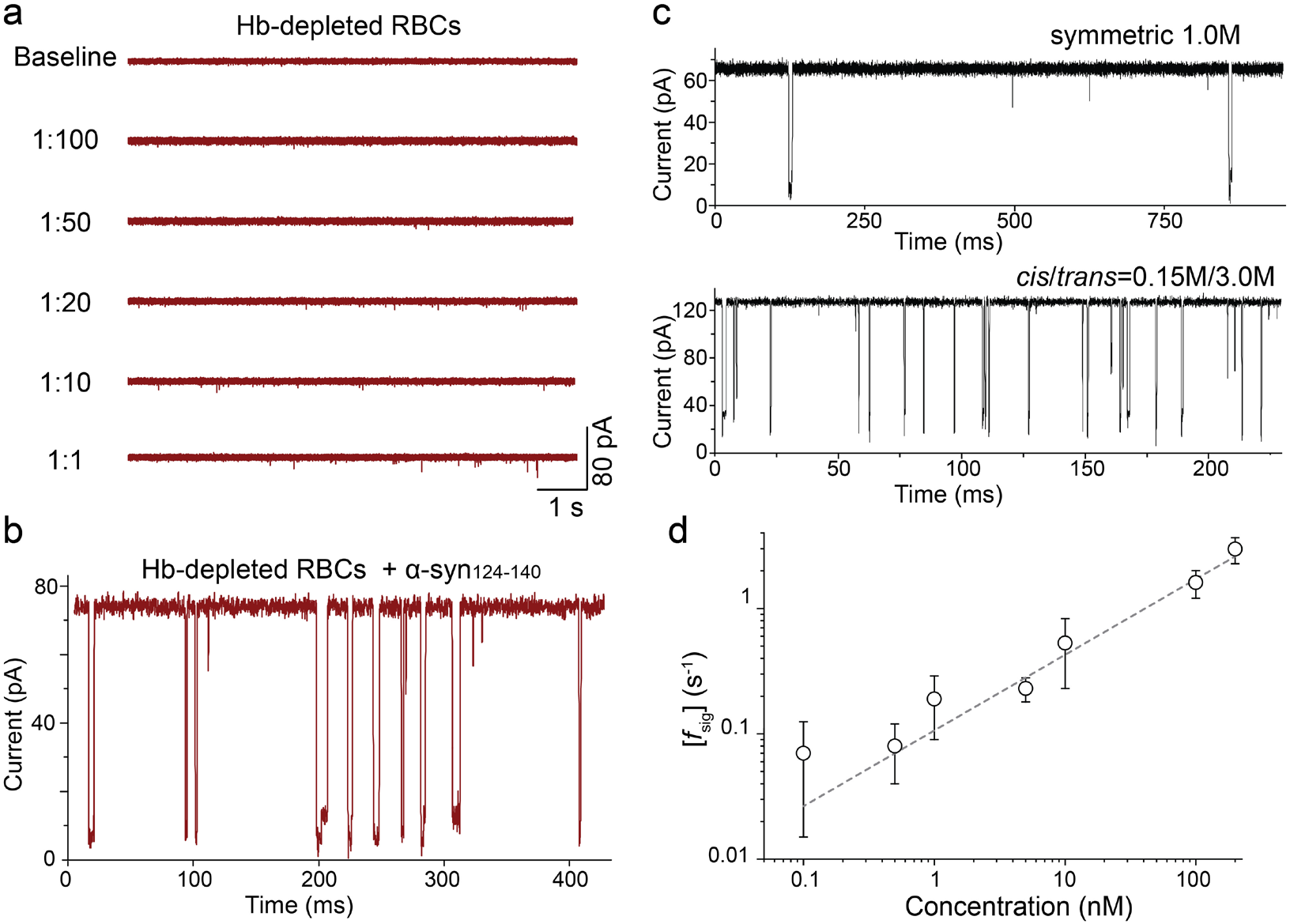
Detection of α-synuclein peptides at low abundance concentration. **(a)** Raw current traces after addition of serial dilutions of Hb-depleted RBC samples. **(b)** Raw current traces of Hb-depleted RBCs sample in presence of 6 µM wt α-syn_124-140_ peptides. The RBCs used here is 1:10 dilution. **(c)** Raw current traces showing signals of α-syn_124-140_ in symmetric salt (top, 1.0 M KCl) and in a gradient salt concentration (bottom, 0.15 M/3.0 M (*cis*/*trans*) KCl). **(d)** The correlation between signal frequency *f*_*sig*_ and the concentration of wt α-syn_124-140_ in gradient salt conditions ranging from 100 pM to 200 nM peptide in the *cis* chamber under 100 mV applied voltage. The error bars represent standard deviations from at least three independent experiments in the same conditions.

When the same PTM occurs at different positions in the peptide sequence, the ionic current and dwell time were modulated differently. For example, the dwell time of pS129 (3.62 ± 0.2 ms) is 6.5 times longer than pY125 (0.55 ± 0.08 ms). We hypothesize that phosphorylation at Y125 (pY125) disrupts the hydrophobic cluster of α-syn_124-140_^47^, all amino acids become more easily exposed and translocate in a linear fashion. In this case, the additional phosphate group of pY125 contributes to the faster translocation under the applied voltages. Similar results were obtained when we compared nY125 and nY136 α-syn_124-140_ (**Supplementary Figure 4a-b**). Unlike nY125, which only showed one population, the relative current of nY136 was less pronounced compared to wt and showed two populations like wt α-syn_124-140_. This suggests that disrupting the local structure facilitates the detection of PTMs in proteins by the aerolysin pore. As reported recently, this could be also achieved by using chemical denaturants such as urea or guanidinium chloride^50^.

**Figure 4.**
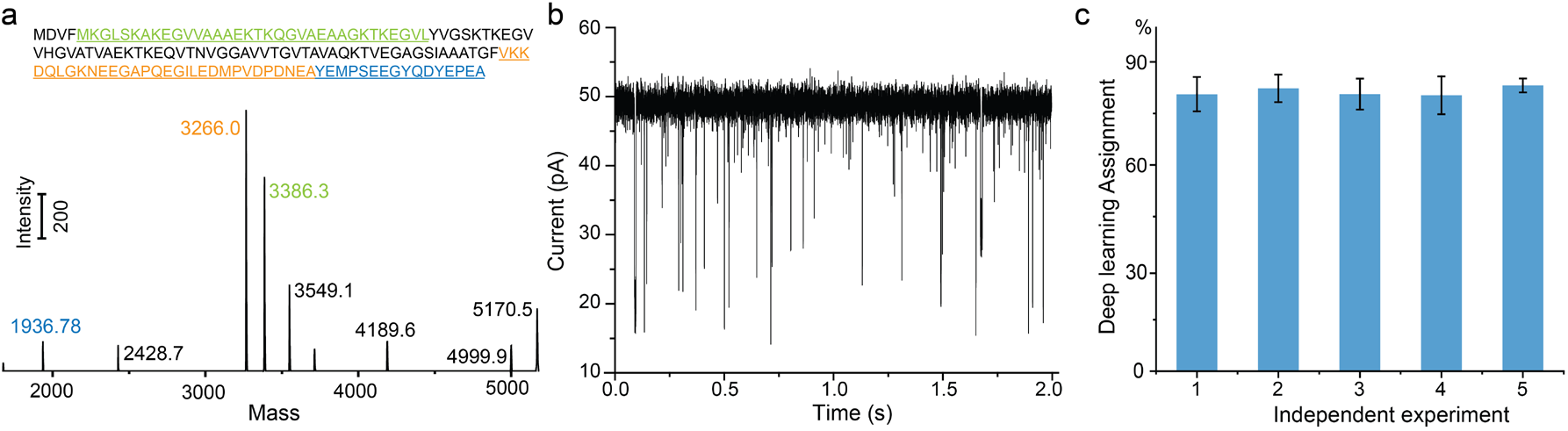
Detection of C-terminal peptides of α-synuclein after CtsD digestion. **(a)** Possible peptide fragments from CtsD digestion based on previous studies_66_ and MS characterization of CtsD digestion of full-length α-synuclein. **(b)** A raw current trace from nanopore measurement of CtsD-digested samples. **(c)** Deep learning result from 5 independent experiments aimed at classifying α-syn_125-140_ from the mixture sample.

To explore the possibility of using the nanopore for the detection of multiple PTMs occurring simultaneously on the same peptide, we measured the ionic current response of double-phosphorylated α-syn_124-140_, pY125pS129, and triple nitrated α-syn_124-140_, nY125nY133nY136 (**Supplementary Figure 4c-d**). Here, the nanopore results showed that compared to the unmodified peptide (wt) and the singly phosphorylated peptides (*i*.*e*., pY125 and pS129), the translocation speed of pY125pS129 (0.45 ± 0.02 ms) was even faster, likely due to its increased negative charge. While the dwell time of nY125nY133nY136 was around 2-fold longer (5.18 ± 0.23 ms) compared to wt and nY136, and slightly longer than nY125: its relative current was the lowest among all peptides (7.2 ± 1.0) since the three modifications contribute to increasing the overall volume of the peptide and therefore induce a deeper blockade of ionic current. In the case of the peptides containing two types of PTMs at different positions, i.e., nitration at Y125 and phosphorylation at S129 (nY125pS129, **Supplementary Figure 4e**), only one population was observed. The relative current of nY125pS129 was between the values of the single modifications, nY125 and pS129, as observed also for the width of the relative current distribution. The dwell time of nY125pS129 was slightly longer than pS129, but identical to nY125. For all these peptides, the dwell time decreases as the voltage increases, indicating that the collected signals are indeed induced by peptides translocating through the nanopore. Altogether, these observations demonstrate that the engineered K238A aerolysin is able to capture the diversity of PTMs.

To classify these PTMs in a more precise, unbiased and automatic way that would allow a translation to clinical applications in the future, we developed a tailored deep learning approach for processing the nanopore current readouts. A long short-term memory (LSTM) recurrent neural network was integrated to read the local extrema of events followed by a multilayer perceptron (MLP) to classify the peptides (**Figure 2b**), a pipeline which is similar to our previous analysis of informational polymers ^51^ (see details in Materials and Methods). We first detected the events with a cutoff threshold at 3σ from the open pore current, and then computed statistical features (i.e., mean residual current, standard deviation and dwell time) and extracted the local extrema of the events **(Supplementary Figure 5** and methods). For each single peptide, we randomly used 75% of the recorded data to feed the deep learning algorithm and train the model; the remaining 25% was used as validation set to compute the accuracy of the model for each single peptide. The events are filtered using a threshold on the predictions confidence of the model which is translated into a selection percentage.

As shown in **Figure 2c**, this approach allows for differentiating wt α-syn_124-140_ from peptides that contain different PTMs at the same position Y125 (i.e., pY125, nY125) with an accuracy of 94% at a 50% selection percentage. Control experiments with a mixture of wt and pY125 at different ratios of concentration were performed to further test the deep learning model (**Figure 2d**). First, a mixture of an equimolar ratio (1:1) was measured in nanopore experiments and the percentage of assignment of wt and pY125 was 52.8 ± 3.5% and 43.2 ± 3.1%, respectively. This is in line with the theoretical predictions. When the ratio of wt:pY125 was changed to 1:3, the percentage of wt assignment decreased to 19.0 ± 0.3% while pY125 increased to 78.2 ± 0.7%. Finally, a mixture ratio of 3:2 (wt:pY125) was tested. As shown in **Figure 2d**, the assignment of wt and pY125 was as expected 61.5 ± 0.1% and 35.0 ± 0.2%, respectively. Therefore, the expectation is that any sequence in the library of peptides (wt, pY125, or nY125) can be identified directly with a confidence level of 94%. Furthermore, mixture experiments of wt and pS129 were performed **(Supplementary Figure 6)**, for which the deep learning model could predict the different mixture ratios with a confidence level of 96%.

Data from wt and the 7 tested α-syn_124-140_ PTM variants (i.e., pY125, nY125, pS129, nY136, pY125pS129, nY125nY133nY136 and nY125pS129) have been used as a training set for the same deep learning model, with a resulting accuracy of 78.2% using a selection percentage of 25% (*i*.*e*., accuracy dependence on selection percentage as shown in **Figure 2e**). If the event is too noisy or doesn’t contain enough information as in the case of short events, the prediction confidence would be lower. If the event is too noisy or doesn’t contain enough information as in the case of short events, the prediction confidence would be lower. The selection percentage allows for a trade-off between accuracy and filtering without requiring heavy filtering during the preprocessing of the events. As illustrated in **Figure 2f**, 13% of pY125 was confused with pY25pS129, because the signals of both pY125 and pY25pS129 showed very short dwell times. This could be further improved if higher bandwidth instruments have been employed. Additionally, 6% of nY125 was confused with nY125nY133nY136, while the other 16% of this peptide was confused with nY125pS129, 10% of nY136 was confused with pS129 and the other 9% was confused with wt. We think this prediction confusion was caused by the similar dwell time between these modified peptides. Nonetheless, the deep learning approach we developed here provides high reading accuracy for all the tested PTMs peptides, particularly for pS129 which is one of the most relevant biomarkers for synucleinopathies. This is encouraging also because a similar deep learning model can be trained for other systems, finding thus general applicability for signal processing in nanopore sensing experiments.

### Detection of α-synuclein peptides at low abundance concentration

To date, most nanopore sensing applications are performed using purified peptides or protein samples (as done hereabove). There is however the need to develop methodologies whereby detection can be performed on biological samples^52^. Nevertheless, even though some nanopores have been used for the detection of biomolecules in biological fluids samples (*e*.*g*., plasma^53,54^, serum^52,55–59^, sweat^60^, urine^60^ and saliva^60^), the task of measuring biomarker peptides is often challenging due to the intrinsic low sensitivity and selectivity of the techniques applied to bodily fluids.

We previously showed that aerolysin pores possess high sensitivity for biomolecular sensing, especially its engineered variants^41,61^. Therefore, we explored if K238A aerolysin was able to detect α-synuclein peptides in clinically relevant conditions, red blood cells (RBCs) devoid of hemoglobin (Hb) proteins. We first determined the dilution necessary to ensure a stable baseline of the ionic current through serial dilutions of Hb-depleted RBC samples. In the literature, no particular instabilities of the lipid bilayer were observed when testing RBCs except when dilutions were lower than 1:50^60^. Based on these previous reports, we scanned a range of dilutions up to 1:100 (**Figure 3a**) in our setting and observed that after the addition of 3 μl of Hb-depleted RBCs extracts into the *cis* chamber no obvious effects were detected for the membrane stability. Interestingly, during the experiments with Hb-depleted RBCs, we could continually collect data for up to three hours, indicating that the aerolysin pore remained invariantly intact likely due to the ultra-stability provided by its unique double β-barrel structure^33^. Moreover, when the concentration of Hb-depleted RBCs samples increased in the chamber, occasional signals were also observed (**Supplementary Figure 7**). Notably, all raw current traces shown in **Figure 3** were collected 30 min after samples were added.

Next, we sought to quantify the presence of α-synuclein wt peptide within the Hb-depleted RBCs, with the aim of mimicking the real complexity of pathological conditions within a clinical setting. Based on the previous tests, dilutions of 1:10 was chosen as the background for Hb-depleted RBCs extracts. As illustrated in **Figure 3b**, after the addition of wt α-syn_124-140_, typical two-level signals were recorded.

While these results represent the potential of aerolysin pores to detect C-terminal α-synuclein peptides directly from an actual clinical sample, another prerequisite towards the development of aerolysin pores into an efficient single-molecule proteomic device for biomarkers detection is the ability to detect molecules in low abundance, as often found in clinical samples. In previous studies, the frequency of signals has been used to quantify the detected molecule. Therefore, we used the equation *f*_sig_ = *k*_on_ *[PTMs]*_*0*_ to explore the theoretical detection limit of aerolysin nanopores for sensing α-synuclein peptides. As illustrated in **Supplementary Figure 8**, we verified that the frequency of blockade events, *f*_sig,_ was proportional to the concentration of α-synuclein ranging from 120 nM to 24 μM, when 100 mV were applied using a symmetric buffer of 1.0 M KCl. The physiological concentration of α-synuclein in RBCs is however much lower than these tested conditions, namely 26.2 ± 3.0 µg/ml^62,63^, which corresponds to 35 nM if 3µl of the original sample was added into the nanopore system. Therefore, to enhance the detection limit of our setting, we measured peptides in a gradient of salt concentration, which was shown to be an efficient way to increase the capture rate for ssDNA^53,64^. An asymmetrical KCl buffer solution consisting of 0.15 M in the *cis* chamber and 3.0 M in the *trans* was thus used (**Figure 3c**), which enabled the detection of wt α-synuclein peptides at far lower concentrations (*i*.*e*., ∼100 pM, **Figure 3d**) than when using canonical symmetric buffer conditions.

Altogether, these results demonstrate that K238A aerolysin nanopores can detect C-terminal peptides of α-synuclein at ∼nM concentration (factoring in the dilution conditions), condition which is overlapping with the concentration at which they are found in a RBC clinical sample. Considering that an even lower detection limit could be achieved by applying higher voltages, minimizing the volume of the chambers or optimizing the pore variants, the possibility to develop an effective nanopore-based tool for the diagnosis of synucleinopathies appears within reach.

### Detection of α-synuclein peptides in complex mixtures

Next, we sought to determine if the presence of multiple fragments, produced for instance by proteolytic cleavage of α-synuclein could influence the nanopore detection of C-terminal peptides bearing the desired PTMs. This aims at mimicking and describing the actual α-synuclein mixtures one could find in an actual clinical sample. For this purpose, we choose to work with cathepsin D (CtsD) protease as previous studies showed that CtsD-mediated digestion of α-synuclein results in the generation of a C-terminal fragment spanning residues 125-140 (α-syn_125-140_), in addition to four other α-synuclein fragments of various length^65,66^ (**Figure 4a**). The C-terminal fragment contains the great majority known disease-associated PTM sites (p/nY125, pS129, nY133, nY136), including all the ones we investigate above for α-syn_124-140_. To the best of our knowledge, CtsD is the only enzyme that cleaves α-synuclein and results in the generation of a peptide fragment that contains all the most relevant PTM sites, thus making it the ideal choice for future studies to detect α-synuclein C-terminal peptides bearing single or multiple PTMs. Other enzymes which cleave within the C-terminal domain, e.g., Glu-C, give rise to much shorter peptides, thus precluding the detection of single peptides containing multiple PTMs^42^.

We incubated CtsD with full-length α-synuclein (see details in Methods) and detected the resulting products by nanopore experiments afterwards. In parallel, the same sample was characterized by MS and as shown in **Figure 4a**, CtsD digestion produced three main peptide fragments, α-syn5-38, α-syn95-124, and α-syn_125-140_. After adding the CtsD-digested α-synuclein sample into the nanopore system, diverse current signals were obtained (**Figure 4b**). Under these conditions, the deep learning model was able to identify the C-terminal peptides α-syn_125-140_ with an average accuracy of 81.4 ± 1.1% from five independent experiments (**Figure 4c**), implying that the other proteolytic fragments, while having an effect of the current readout do not compromise the ability to detect the α-syn_125-140_ fragment. It should be noted that there is one amino acid difference between peptides used to train the deep learning model and this α-syn_125-140_ fragment; difference which however does not seem to affect significantly the accuracy of predictions, likely because Ala124 does not contribute in a significant way to modulate the current signal during translocation, while not holding any PTM. Moreover, as α-syn_5-38_ is positively charged, we do not expect it to be captured by the nanopore when using +100 mV voltage. Therefore, the only possible fragment that may interfere with the measurement is α-syn_95-124_ since it is negatively charged. However, it should be noted that the length of α-syn_95-124_ is two times longer than α-syn_125-140_ which may lead to a low capture rate in the conditions used here. As shown for DNA, the capture of long ssDNA is dramatically reduced in the aerolysin nanopore^67^.

## Discussion

In this work, we have shown that an engineered aerolysin nanopore can detect and distinguish peptides carrying different types and number of PTMs occurring at different residues. In addition, the measurements can capture subtle structural features that are challenging to be characterized by other biophysical methods without fluorescently labeling or modifying the peptides/proteins. Using a deep learning model for signal processing, all investigated PTMs could be automatically identified in a supervised context, which means this approach can be scaled up to identify more PTMs or scaled down to fit a specific application. Importantly, this nanopore approach can reach a detection limit as low as 100 picomolar concentration and is amenable for high-throughput applications, which are challenging for other techniques such as MS. Finally, one major advantage of this nanopore-based approach is that it enables simultaneous detection of several protein PTMs which is difficult to achieve using immunoassay/antibody-based methods. This is because the presence of multiple PTMs alters the biochemical properties of the antibody-targeting epitopes.

Altogether, our findings open promising avenues for developing biological nanopores into efficient single-molecule proteomic devices and diagnostic tools. We envision that this could be achieved by detecting the circulating peptides directly, or by isolating target proteins from various biological fluids using specific antibodies^68^ and then digesting them into smaller peptides and subsequently detecting them through a nanopore as reported in the recent studies^65,69^. In the future, clinical samples could be treated directly with the specific protease, rather than extracting the target proteins, and subsequently detected and classified using nanopores. Several very recent studies have shown that nanopore sensors can be used identify different types of proteins by assessing characteristic blockage event features linked to peptide fragments produced by digestion of the proteins with proteases^70,71^. Such nanopore-based molecular diagnosis platform would hold also promise to detect multiple biomarkers simultaneously, broadly extending the potential field of application, for instance for the detection of biomarker signatures for cancer diagnosis^72^.

The ability of the aerolysin pore to detect and distinguish between long peptides bearing multiple PTMs also paves the way for more precise remapping of PTMs in proteins at the single-molecule level, which remains a challenge for MS and other methods. In conclusion, the nanopore-based technology presented here, besides the natural advantages of being fast, cheap, label-free, and high-throughput, provides the possibility to be developed into a portable diagnostic device with medical and commercial potential.

## Materials and Methods

### Synthesis of C-terminal α-synuclein peptides and their PTMs

The majority of peptides used in nanopore experiments were produced as described in previous works^44–46^. A few peptides (oM127 and wt showed in Supplementary Figure 2) were provided by GenicBio Limited. All peptides were characterized by Liquid chromatography-mass spectrometry (LC-MS) as previously described^68^. Their purity was also assessed by UPLC analysis, on a Waters Acquity H-Class system using a C18 column (with UV detection at 214 nm and run time of 4 min (gradient 10% to 90% acetonitrile) with 0.6 mL/min flow rate.

RBCs were prepared similarly as previously described^73^. Briefly, EBCs were lysed and further treated using the Hemovoid kit (Biotech Support Group), aiming to remove hemoglobin (the most abundant protein in RBCs) but also to enrich low abundant proteins such as α-synuclein. Importantly, all biological fluids used in this study are derived from healthy controls.

CtsD digestion experiments were performed in a total volume of 100 µl, 2 µl of 500 µM CtsD (Sigma-Aldrich Chemie GmbH, Buchs, SG Switzerland), 3 µl of 100µM full-length α-synuclein and 95 µl buffer (40mM sodium acetate, 50mM NaCl and 5mM DTT, pH 5) incubated at 37°C for 20h at 300 rpm in a ThermoMixer C (Eppendorf).

### Aerolysin productions

The recombinant K238A aerolysin proteins were generated from the aerolysin gene in the pET22b vector with a C-terminal hexa-histidine tag as described in our previous work^41,51^, and then expressed and purified from BL21 DE3 pLys *E. coli* cells. Cells were grown to an optical density of 0.6-0.7 in Luria-Bertani (LB) media. Protein expression was induced by the addition of 1 mM isopropyl β-D-1-thiogalactopyranoside (IPTG) and subsequent growth overnight at 20α C. Cell pellets were resuspended in lysis buffer (20 mM Sodium phosphate pH 7.4, 500 mM NaCl) mixed with cOmplete™ Protease Inhibitor Cocktail (Roche) and then lysed by sonication. The resulting suspensions were centrifuged (12.000 rpm for 35 min at 4α C) and the supernatants were purified through a HisTrap HP column (GE Healthcare) previously equilibrated with lysis buffer. The protein was eluted with a gradient over 40 column volumes of elution buffer (20 mM Sodium phosphate pH 7.4, 500 mM NaCl, 500 mM Imidazole), and buffer exchanged into a final buffer (20 mM Tris, pH 7.4, 500 mM NaCl) using a HiPrep Desalting column (GE Healthcare). The purified protein was flash-frozen in liquid nitrogen and stored at -20α C.

### Single-channel recording experiments

Phospholipids of 1,2-Diphytanoyl-*sn*-glycero-3-phosphocholine (DPhPC) powder (Avanti Polar Lipids, Alabaster, USA) were dissolved in octane (Sigma-Aldrich Chemie GmbH, Buchs, Switzerland) for a final concentration of 1.0 mg per 100 μl. Purified protein was diluted to the concentration of 0.2 μg/ml and then incubated with Trypsin-agarose (Sigma-Aldrich Chemie GmbH, Buchs, SG Switzerland) for 2 h under 4°C temperature. The solution was finally centrifuged to remove trypsin.

Nanopore single-channel recording experiments were performed on Orbit Mini equipment (Nanion, Munich, Germany) and an Axopatch 200B Amplifier system (Molecular Devices, San Jose, USA). In the Orbit Mini set-up, DPhPC membranes were formed across a MECA 4 recording chip that contains a 2 x 2 array of cylindrical 50 µm diameter in a highly inert polymer. Each of the four cavities contains an individual integrated Ag/AgCl-microelectrode and sustains one DPhPC bilayer. If not indicated otherwise, the measurement chamber temperature was set to 20°C. Data was collected at 10 kHz sampling rate with a 5 kHz low-pass filter.

Ionic strength gradient experiments were carried out as follows. Teflon films with 50 µm apertures were mounted in Teflon chambers using high-vacuum grease (Dow Corning Corporation, Midland, MI, USA). The films separated two compartments (cis/trans) only connected through the Teflon film aperture, with one Ag/AgCl electrode in each compartment. Apertures were pretreated with 1 µL 2 % (v/v) hexadecane in hexane on both sides using a standard pipette and the chamber was mounted in the recording setup. DPhPC bilayers were formed by folding as described previously^74,75^. Briefly, electrolyte solution was added to both sides taking care that the level stayed below the aperture, lipids (10 mg/mL in pentane) were added onto the electrolyte surface in both compartments. After the pentane evaporated, the electrolyte level was raised above the aperture and a lipid bilayer was formed. The quality of the lipid bilayer was monitored through its capacitance and its stability was verified through the application of 150 mV over the course of at least 5 minutes. After peptide addition, the cis chamber was carefully mixed by pipetting up and down. Currents were sampled at 200 kHz and low-pass filtered at 100 kHz with the Axopatch 200B (Molecular Devices, LLC., San Jose, CA, USA).

Peptides (lyophilized powder) were pre-diluted in 10 mM Tris and 1.0 mM EDTA solution (pH=7.4) to a stock concentration of 500 µM and added to the *cis* side of the chamber in 1.0 M KCl solution buffered with 10 mM Tris and 1.0 mM EDTA (pH=7.4) to the final concentration indicated in the figure caption. All experiments shown here were repeated with at least 4 different pores.

### Signals processing and classifications using deep learning

The signal processing was done in the same way as previously reported^51^. The open pore current distribution is measured by fitting a Gaussian function on the peak distribution of current with the highest mean current. The events are extracted using a current threshold at 3σ from the open pore current distribution. The relative current percentage (*I/I*_*0*_) is computed from the mean open pore current (*I*_*0*_) and the mean residual current (*I*). The dwell time, average relative current, relative standard deviation of the current 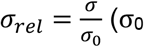 is the value of the open pore current standard deviation and σ is the residual current standard deviation) and local extrema are computed. The events are selected on the basis of the dwell time (0.2 to 100.0 ms) and the average relative current (0 to 40%) discarding the events that are too short, too long or that do not block the current sufficiently.

The machine learning pipeline is composed of two steps. The first one is the classification of every event and the second is the assessment of the quality of the prediction of the classifier. The neural network architecture for both the classification and the assessment is a long short-term memory (LSTM) neural network followed by a multilayer perceptron (MLP) using the position in time and relative current of the local extrema for each event as input features. The features are rescaled by a fixed factor to decrease the training time. The classifier is composed of a LSTM with state size 128 without any activation function followed by 6 fully connected hidden layers of size 256 with rectified linear unit (ReLU) as activation functions and finally an output layer of size 8 with softmax activation function. The assessment is done with a scaled down version of the classifier with a LSTM with a state of size 32, 3 fully connected hidden layers of size 64 with hyperbolic tangent activation functions and an output layer of size 1 with sigmoid activation function. The neural networks for the classification and assessment are trained together using a 3-parts loss function. The first part is the full classification cross-entropy loss of the predictions from the classifier and the peptides label. The second part is the assessment of cross-entropy loss between the predicted and actual prediction validity from the classifier. The third part is the reinforcement classification loss which is the full classification cross-entropy loss scaled by the assessment prediction.

## Acknowledgments

This research was supported by the Swiss National Science Foundation (to M.D.P. and PR00P3_193090 to C.C), the European Union’s Horizon 2020 Research and innovation program under the Marie Skłodowska-Curie grant agreement No. 665667, Peter and Traudl Engelhorn Foundation and Synapsis foundation (2019-CDA02 to C.C). The authors also thank M. J. Marcaida for support with aerolysin production and R. Kolla for the discussion on the peptides’ characterization.

## Author Contributions

C.C., M.D.P. and H.L. conceived the project and designed the study. C.C. performed the experiments on Orbit Mini, supervised all single-channel experiments and did data analysis. P.M. characterized all C-terminal α-synuclein peptides and prepared all clinical samples. L.F.K. performed signal processing and developed the deep learning pipeline.

J.F.B.J conducted CtsD digestion and nanopore experiments. S.F.M. and V.R. carried out experiments on the asymmetric buffer. A.C. synthesized C-terminal α-synuclein peptides with different PTMs forms. C.C., L.F.K., P.M, S.F.M., H.L., and M.D.P. interpreted the data. C.C., H.L., and M.D.P. wrote the manuscript with input from all authors.

