## Supplemental Figures S1-S8 for "Deep learning-assisted single-molecule detection of protein post-translational modifications with a biological nanopore"

**This PDF file includes:**

Figures S1 to S8

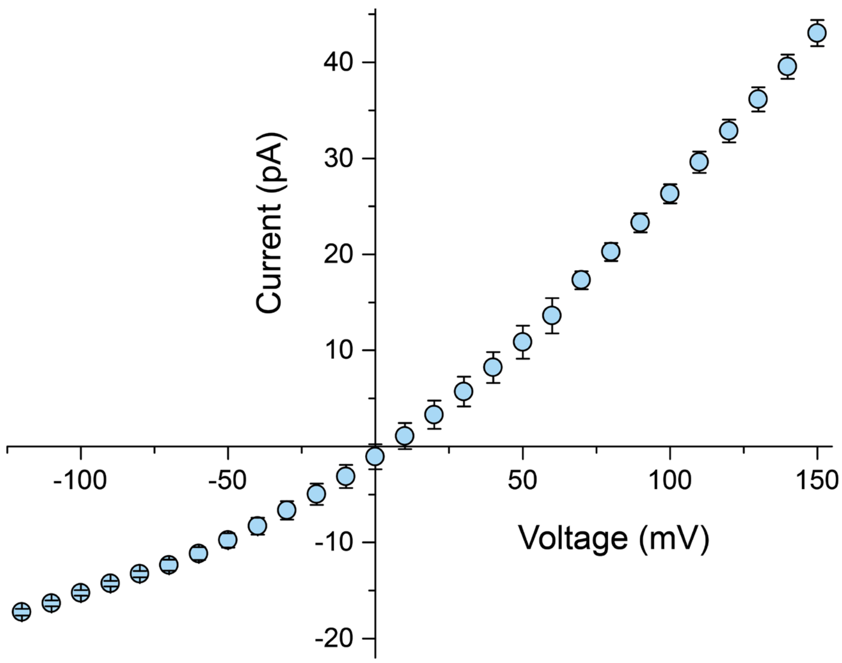


**Supplementary Figure 1 |** IV curve for K238A engineered aerolysin nanopore in a buffer of 8.0 M urea, 1.0 M KCl, 10 mM Tris, 1.0 mM EDTA, pH 7.4. This engineered aerolysin pore showed a folded structure and function well in the buffer of 8.0 M urea solution.

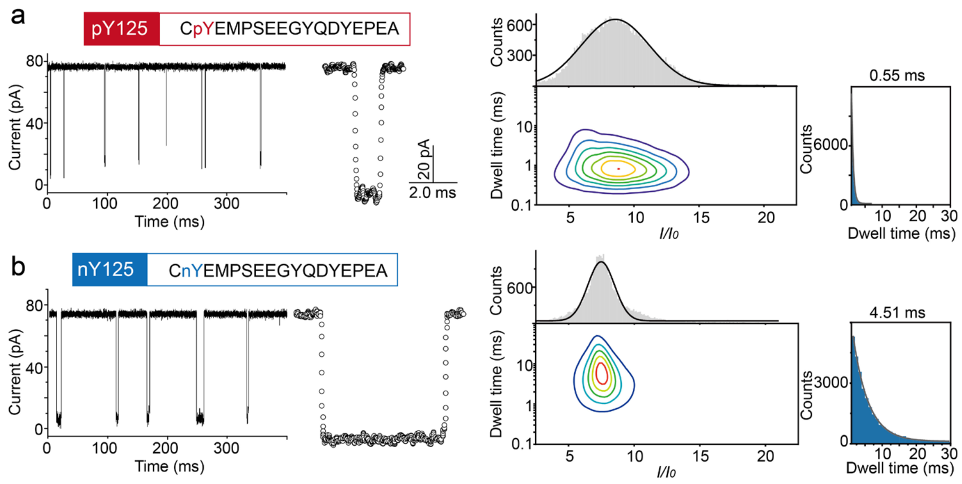


**Supplementary Figure 2 | Ionic current signals of different PTM types at the same position.** Sequence, raw current trace, typical event, contour plot, *I/I₀* percentage and dwell time histograms of α-syn$_{124\text{-}140}$ peptides for **(a)** pY125 and **(b)** nY125.

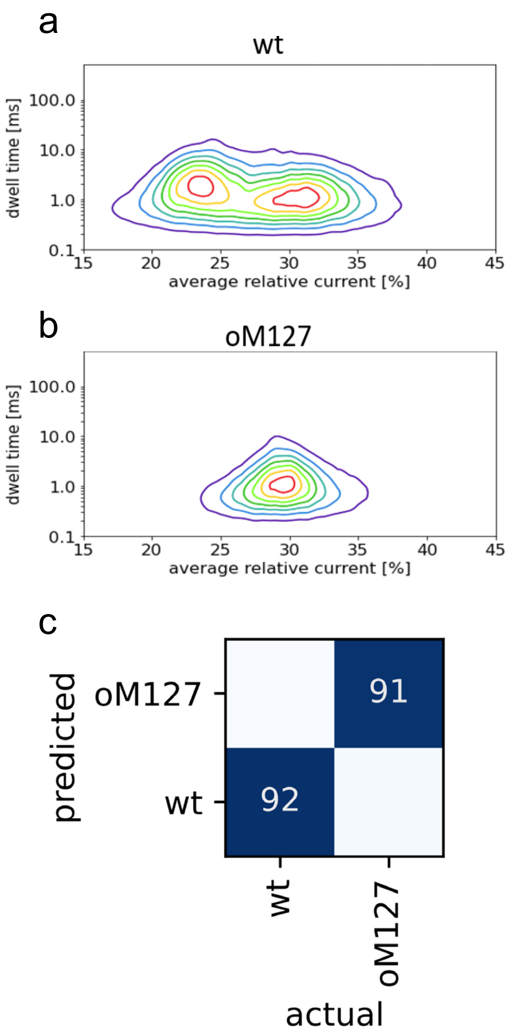


**Supplementary Figure 3 |** Contour plot for **(a)** wt α-syn$_{124\text{-}140}$ and **(b)** oM127 peptides. **(c)** Confusion matrix for wt and oM127 classification. Columns represent actual peptides from the test set, while rows are the peptides that the deep learning algorithm assigned them to. Peptides were provided by GenicBio Limited.

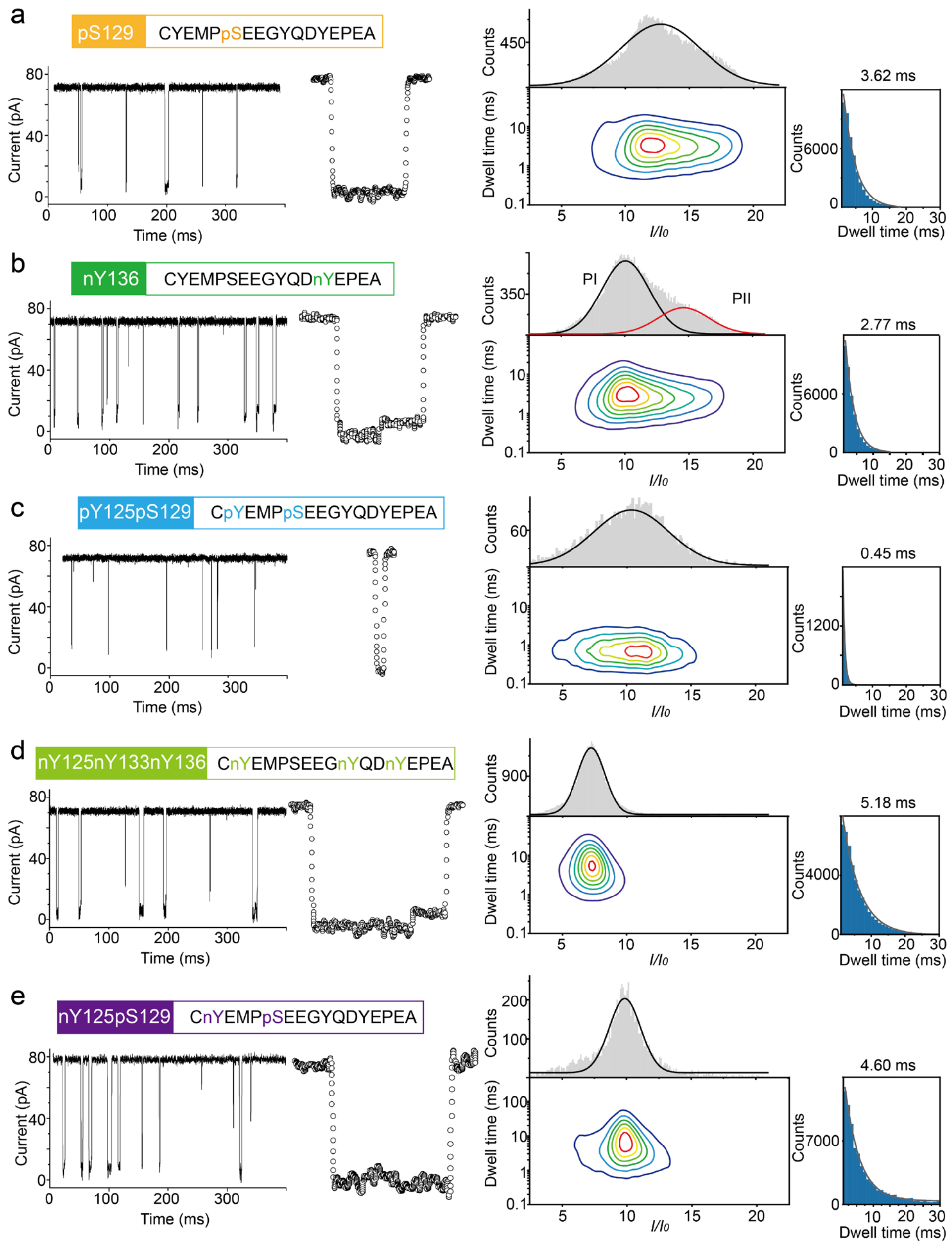


**Supplementary Figure 4 | Ionic current signals of same PTM type at different and multiple positions, and combination of different PTMs.** Sequence, raw current trace, typical event, contour plot, $I/I_0$ percentage and dwell time histograms of α-syn$_{124\text{-}140}$ peptides for **(a)** pS129, **(b)** nY136, **(c)** pY125pS129, **(d)** nY125nY133nY136 and **(e)** nY125pS129.

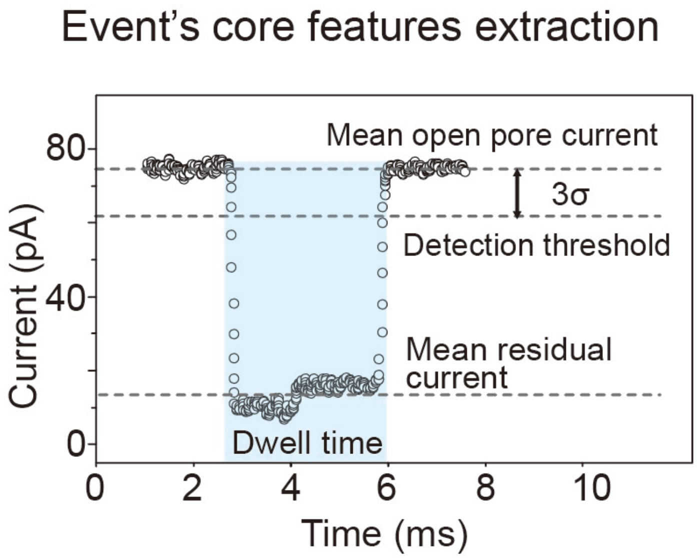


**Supplementary Figure 5 I** Illustration of event's features extraction.

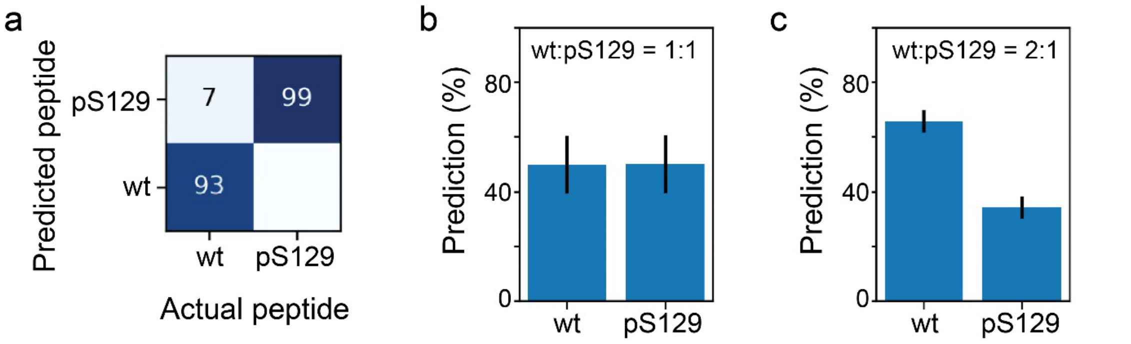


**Supplementary Figure 6 | Mixture experiments of wt and pS129**. **(a)** Confusion matrix of wt and pS129 peptides obtained by deep learning approach. **(b)** Prediction of the mixture experiment of wt and pS129 at a concentration ratio of 1:1. The prediction for wt and pS129 is 49.9 ± 10.6% and 50.1 ± 10.6%, respectively. This gave a predicted ratio of WT/pS129 = 0.995 ± 0.417, which is very close to 1, validating the ML model we developed. **(c)** Prediction of the mixture experiment of wt and pS129 at a concentration ratio of 2:1. the prediction of wt increase to 65.7 ± 3.4% while pS125 decrease to 34.3 ± 3.4%, leading to a predicted ratio of WT/pS129 = 1.916 ± 0.288, which is in line with the mixture ratio.

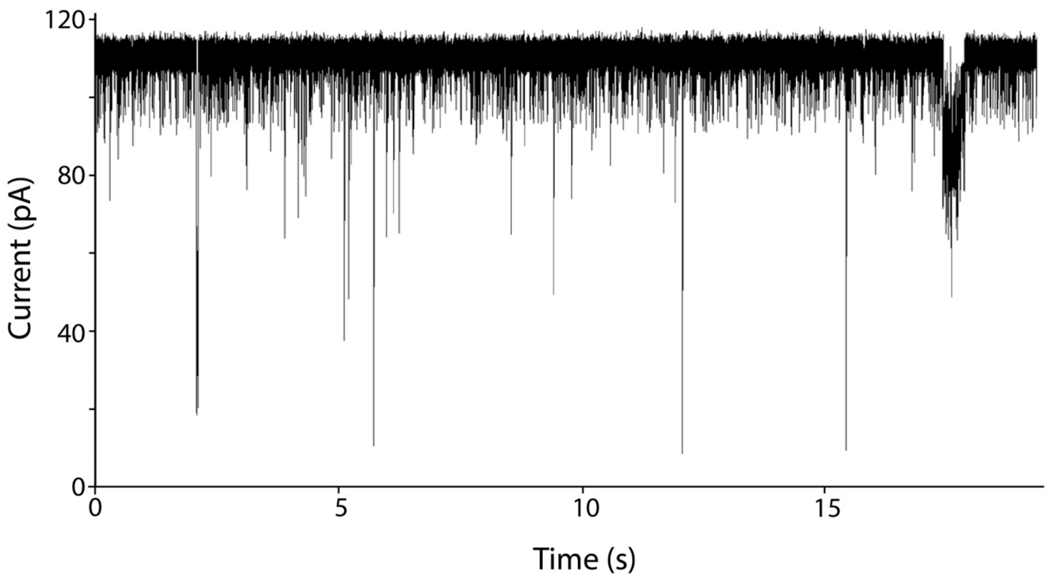


**Supplementary Figure 7 |** Typical raw current trace for a nanopore measurement of red blood cell lysates devoid of hemoglobin after 3 hours' recording.

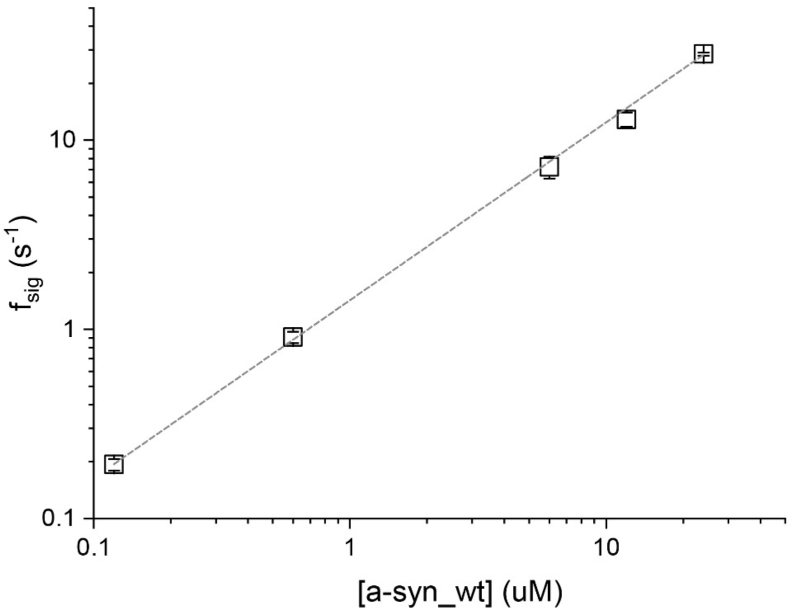


**Supplementary Figure 8 |** Correlation between signal frequency $f_{sig}$ and the concentration of wt α-synuclein ranging from 120 nM to 24 µM, which was calculated by applying 100 mV in a symmetric 1.0 M KCl buffer.
